# Helminth parasites decrease survival probability in young red deer

**DOI:** 10.1101/2022.03.03.482858

**Authors:** Claudia I. Acerini, Sean Morris, Alison Morris, Fiona Kenyon, David McBean, Josephine M. Pemberton, Gregory F. Albery

## Abstract

Helminths are common parasites of wild ungulates that can have substantial costs for growth, mortality, and reproduction. While these costs are relatively well documented for mature animals, knowledge of helminths’ impacts on juveniles is more limited. Identifying these effects is important because young individuals are often heavily infected, and juvenile mortality is an important process regulating wild populations. Here, we investigated associations between helminth infection and overwinter survival in juvenile wild red deer (*Cervus elaphus*) on the Isle of Rum, Scotland. We collected faecal samples non-invasively from known individuals and used them to count propagules of three helminth taxa (strongyle nematodes, *Fasciola hepatica*, and *Elaphostrongylus cervi*). Using generalised linear models, we investigated associations between parasite counts and overwinter survival for calves and yearlings. Strongyles were associated with reduced survival in both age class, and *F. hepatica* was associated with reduced survival in yearlings, while *E. cervi* infection showed no association with survival in either age class. This study provides observational evidence for fitness costs of helminth infection in juveniles of a wild mammal, and suggests that these parasites could play a role in regulating population dynamics.

## 1. Introduction

Parasites are ubiquitous in natural populations and are often costly to the hosts they infect (Hudson *et al*., 2002). Whilst the consequences of parasitism in mammals are well documented for domestic livestock, evidence of their effects in wild populations is far more limited due to the practical difficulties of collecting long-term parasitological data from wild hosts – particularly large, long-lived mammals (Coulson *et al*., 2018; Wilson *et al*., 2003). Wild mammals are typically infected with gastrointestinal helminth parasites, a paraphyletic clade of macro-parasitic worms, including tapeworms (Cestoda), roundworms (Nematoda), and flukes (Trematoda) (Taylor *et al*., 2015b). These parasites display a variety of life histories and induced pathologies in their hosts (McSorley and Maizels, 2012). Most frequently, helminths invade their host via the gastrointestinal tract, after free-living larval stages are consumed by the host (Taylor *et al*., 2015b). Adult helminths live, feed, and reproduce within their hosts and their propagules are excreted into the environment with the faeces, from which they spread to other hosts either directly or indirectly, via an intermediate host (Taylor *et al*., 2015b). Quantification of infection is possible by counting these propagules within a host’s faeces in a method known as faecal egg counts (FECs) (Taylor *et al*., 2015a). This non-invasive measure can be used as a proxy for an individual’s parasite burden, defined as the actual quantity of adult helminths within the host (Budischak *et al*., 2015). Parasite count often varies with both extrinsic and intrinsic host factors. Age-dependent parasitism is common; juveniles are often the most heavily parasitised members in a population, predominantly attributed to their naïve immune systems and prioritisation of resources for growth rather than immunity (Ashby and Bruns, 2018; Wilson *et al*., 2003). Juveniles are a key demographic group and any parasite-mediated effects on their survival could play a role in population regulation (Gaillard *et al*., 2000).

The European red deer (*Cervus elaphus*) is a large ungulate that has great ecological importance as a wide-ranging herbivore and source of livestock diseases (Böhm *et al*., 2007; Fuller and Gill, 2001). Red deer are abundant in Scotland and culling regimes for population regulation have provided the basis for many parasitological investigations. These studies have documented the prevalence of endoparasites in Scottish red deer, including multiple species of strongyle nematodes (a family of worms whose eggs are indistinguishable by microscopy and so grouped together in assays), lungworms (*Dictyocaulus spp*.), the tissue nematode (*Elaphostrongylus cervi*), *Sarcocystis spp*., and the common liver fluke (*Fasciola hepatica*) (Böhm *et al*, 2006; Irvine *et al*., 2006; French *et al*., 2016). The study population of wild red deer on the Isle of Rum provide an excellent system for investigating the fitness consequences of parasitism. Longitudinal individual-based monitoring enables collection of complete life history information, and parasite data from non-invasive faecal sampling (Albery *et al*., 2021). The population hosts a variety of helminth parasites, the most prevalent taxa being strongyle nematodes, *F. hepatica* and *E. cervi* (Albery *et al*., 2018). Juvenile deer tend to be more heavily parasitised than adults, with calves (≤12 months old) showing the highest strongyle intensities and yearlings (13-24 months old) showing the highest *F. hepatica* and *E. cervi* intensities (Albery *et al*., 2018). Mortality rates are high among juveniles, with many of these deaths occurring over the winter months (January-March), when environmental conditions are harshest and food is limited (Clutton-Brock *et al*., 1987; Coulson *et al*., 1997). Juvenile overwinter survival may be influenced by the extent of parasite infection. In other wild ungulates, helminth infection in juveniles has been shown to cause mortality over winter periods, exacerbating the effects of food shortage (Coltman *et al*., 1999). Strongyle infection negatively impacts future reproductive success and survival in adult female deer in the Rum study population (Albery *et al*., 2021), but to date there have been no investigations into the fitness costs of juvenile parasitism in this population.

Here, we investigate associations between survival probability of juvenile red deer on the Isle of Rum, and infection of strongyle nematodes, *F. hepatica* and *E. cervi*, quantified from faecal samples collected at three times of year. We predict that increases in helminth parasite burden in young red deer will decrease their subsequent overwinter survival probability.

## 2. Methods

### 2.1 Data collection

This study uses data collected between 2016 and 2020 from a wild population of red deer situated on the North block of the Isle of Rum, Scotland. A detailed description of the study system and field data collection can be found in Clutton-Brock *et al*. (1982). After many years of study, the deer are relatively habituated to human presence. The ‘deer year’ begins on May 1^st^, marking the start of the calving season (May-July). During this time pregnant female deer are monitored daily for when they give birth to a single calf. Within a few hours of birth, calves are caught, sex determines, weighed and marked with a combination of collars, tags, and ear punches, to allow individual identification throughout their lives. Regular censuses of the population allow accurate individual life history data to be collected.

Faecal samples were collected in spring (April), summer (August) and autumn (November). A detailed description of faecal sampling and parasitological methods can be found in Albery *et al*. (2018). Individually recognised deer were observed defaecating from a distance and the faeces were collected as quickly as possible without disturbing the deer. In each season as many different individuals as possible were sampled. Faecal samples were kept as anaerobic as possible in re-sealable plastic bags and refrigerated at 4°C to prevent the hatching or development of parasite propagules until parasitological analysis was performed (within 3 weeks of collection) (Albery *et al*., 2018).

From a faecal subsample, parasite propagule counts were conducted for strongyle nematodes, *Fasciola hepatica* and *Elaphostrongylus cervi* as detailed in Albery *et al*. (2018). Briefly, strongyle nematode FECs were conducted via a sedimentation-salt flotation method accurate to 1 egg per gram (EPG) (Kenyon *et al*., 2013; Albery *et al*., 2018); faecal samples were homogenised in water to suspend eggs, the suspension was then filtered, centrifuged at 200 x g for 2 minutes, and the supernatant was removed using a vacuum. Retentate was mixed with saturated salt solution and then centrifuged again. The less dense strongyle eggs that floated to the surface were collected and counted under 4X magnification. *F. hepatica* eggs were counted by a sedimentation method (Taylor *et al*., 2015a); faecal matter was homogenised with water and filtered. The sample was then left to sediment; the dense eggs that sank to the bottom were separated from the lighter material above and stained with methylene blue to facilitate counting under 4X magnification. *E. cervi* larvae were counted by a baermannization method (Gajadhar *et al*., 1994); faecal matter was wrapped in muslin cloth, submerged in a tube of water and left for 20-24 hours for the larvae to emerge and fall to the bottom of the tube. The supernatant was then removed, and the remaining larvae counted under 40X magnification. Propagule counts were divided by the mass of the faecal subsample used, to give a measure of parasitic burden as eggs per gram of faecal matter for strongyles and *F. hepatica* (EPG) or larvae per gram of faecal matter for *E. cervi* (LPG). Our analysis uses faecal propagules counts included in Albery *et al*. (2018) collected in 2016, and additional samples collected in 2017, 2018 and 2019.

### 2.2 Statistical analysis

All statistical analysis was performed in R version 4.0.3 (RStudio Team, 2021). For calves and yearlings, we calculated prevalence (%) and mean FEC of strongyles, *F. hepatica* and *E. cervi* in the spring (April), summer (August), and autumn (November). We do not investigate parasite counts for yearlings sampled in the spring (April) as they have already survived over the winter period and so are not informative for survival analysis. We used binomial generalised linear models (GLM) to explore the association of parasitic burden with subsequent overwinter survival in calves and yearlings. Parasite burden, determined by FECs, was log(count +1) transformed in all cases, to approximate normality. To investigate the survival of a calf through their first winter, we conducted generalised linear models with a logit-link function, using faecal sample data from the summer (model A) and autumn (model B) before the calves’ first winter. In both models we included a response variable of first winter survival (binary; survived (1) or died (0)) and explanatory variables of sex (categorical; female, male), sample deer year (categorical) and strongyle count per gram of faeces (continuous). We included *F. hepatica* count per gram of faeces (continuous) as an explanatory variable in model B but not model A, as *F. hepatica* infection is prepatent and FECs are not meaningful when sampled from calves in the summer at the age of two to three months. We did not fit *E. cervi* count in either model as infection is prepatent and FECs are not meaningful when sampled from calves aged up to six months in the summer and autumn (Albery *et al*., 2018; Gajadhar *et al*., 1994). To investigate yearlings’ survival through their second winter, we conducted generalised linear models with a logit-link function, using faecal sample data from the spring (as calves; model C), summer (as yearlings; model D) and autumn (as yearlings; model E) before the individuals’ second winter. In all three models we included a response variable of second winter survival (binary; survived (1) or died (0)) and explanatory variables of sex (categorical; female, male), deer year (categorical), strongyle count per gram of faeces (continuous), *F. hepatica* count per gram of faeces (continuous) and *E. cervi* count per gram of faeces (continuous). Survival was 97.1% for calves and 98.1% for yearlings in deer year 2019, preventing us from fitting a survival model to this year. For this reason, before running the models, we removed samples corresponding to calf and yearling overwinter survival in the deer year 2019; samples taken in the summer and autumn of the same deer year (August and November 2019) and samples taken in spring of the previous deer year (2018; April 2019).

## 3. Results

Strongyle prevalence and mean intensities were higher in calves than yearlings, peaking in calves sampled in the spring aged ten to eleven months. Strongyle prevalence and intensity was lowest in the autumn. *F. hepatica* prevalence and intensity peaked in spring and dropped in the summer and autumn. *E. cervi* showed the highest mean intensity across all parasite taxa, with calves sampled in the spring displaying the highest mean counts. Prevalence of *E. cervi* was highest in yearlings sampled in the autumn (Table 1).

**Table 1.** Prevalence (%) and mean faecal propagule counts of strongyles, *F. hepatica* (in eggs per gram of faeces) and *E. cervi* (in larvae per gram of faeces) in calves and yearling red deer sampled in the spring, summer, and autumn across all years (2016-2019). ‘P’ indicates parasite is prepatent in the sample and so prevalence and mean propagule counts are not meaningful.

| Sample |  | Strongyles |  | *F. hepatica* |  | *E. cervi* |  |
| --- | --- | --- | --- | --- | --- | --- | --- |
|  |  | Prevalence (%) | Mean count [range] (EPG) | Prevalence (%) | Mean count [range] (EPG) | Prevalence (%) | Mean count [range] (LPG) |
| Summer | Calves (N=141) | 87.2 | 31.1 [0-194.0] | P | P | P | P |
|  | Yearlings (N=91) | 89.0 | 14.8 [0-169.0] | 83.5 | 11.5 [0-46.7] | 85.7 | 40.5 [0-249.2] |
| Autumn | Calves (N=159) | 47.2 | 3.91 [0-33.0] | 81.1 | 8.86 [0-114] | P | P |
|  | Yearlings (N=91) | 42.9 | 3.40 [0-45.0] | 83.5 | 11.3 [0-104.5] | 49.3 | 49.3 [0-371.2] |
| Spring | Calves (N=90) | 95.5 | 70.3 [0-468.0] | 90 | 26.3 [0-132.0] | 91.9 | 91.9 [0-817.4] |

A full listing of model effect sizes is displayed in Table 2. Below we provide mean parasite effect sizes for each survival model, on the logistic-link scale as log(parasite count + 1). Overall, 63.1% of calves survived through their first winter and 84% of yearlings survived through their second winter (excluding data corresponding to overwinter survival in deer year 2019, which was not used in survival analysis). Calf and yearling survival models consistently revealed a significant negative association between faecal strongyle count and subsequent winter survival in each sampled season. A calf’s summer strongyle FEC was negatively associated with their first winter survival (model A, -0.513 ±0.193, p=0.008). Calves that had the lowest summer strongyle FECs (0 EPG, 4.6% of samples) had a 90.0% probability of surviving their first winter, whilst calves with the highest summer strongyle FECs (>40 EPG, 32.1% of samples) had a <57.3% probability of survival (Figure 1A). A calf’s autumn strongyle FEC was negatively associated with their first winter survival (model B, -0.858 ±0.300, p=0.004). Calves that had the lowest autumn strongyle FECs (0 EPG, 56.9% of samples) had an 81.3% probability of survival, whilst calves with the highest autumn strongyle FECs (>10 EPG, 17.9% of samples) had a <35.7% probability of survival (Figure 1B).

**Table 2.** Results from binomial generalised linear models predicting calf and yearling overwinter survival using parasite FEC data collected in different seasons prior to winter (as described in table subheadings). Estimates are given on the logistic scale. Negative estimates indicate a reduction in survival probability. Significant effects are given in bold text.

|  | Estimate | Std. Error | Z value | Pr(>\|z\|) |
| --- | --- | --- | --- | --- |
| **Model A (Summer, calf survival, n=109)** |  |  |  |  |
| (Intercept) | 2.316 | 0.790 | 2.931 | **0.003** |
| Log(Strongyles EPG + 1) | -0.513 | 0.193 | -2.654 | **0.008** |
| Sex [male] | -0.158 | 0.426 | -0.372 | 0.710 |
| Deer year [2017] | -0.536 | 0.481 | -1.116 | 0.264 |
| Deer year [2018] | 0.555 | 0.581 | 0.954 | 0.340 |
| **Model B (Autumn, calf survival, n=123)** |  |  |  |  |
| (Intercept) | 0.729 | 0.549 | 1.328 | 0.184 |
| Log(Strongyles EPG + 1) | -0.858 | 0.300 | -2.860 | **0.004** |
| Log(*F. Hepatica* EPG + 1) | -0.002 | 0.181 | -0.010 | 0.992 |
| Sex [male] | -0.502 | 0.414 | -1.212 | 0.225 |
| Deer year [2017] | 1.632 | 0.819 | 1.992 | **0.046** |
| Deer year [2018] | 1.714 | 0.703 | 2.438 | **0.015** |
| **Model C (Spring, yearling survival, n=67)** |  |  |  |  |
| (Intercept) | 3.451 | 2.153 | 1.603 | 0.109 |
| Log(Strongyles EPG + 1) | -0.869 | 0.372 | -2.339 | **0.019** |
| Log(*E. Cervi* LPG + 1) | 0.212 | 0.218 | 0.974 | 0.330 |
| Log(*F. Hepatica* EPG + 1) | -0.086 | 0.296 | -0.291 | 0.771 |
| Sex [male] | 0.478 | 0.731 | 0.654 | 0.513 |
| Deer year [2016] | 0.803 | 0.784 | 1.025 | 0.305 |
| Deer year [2017] | 1.947 | 1.236 | 1.576 | 0.115 |
| **Model D (Summer, yearling survival, n=64)** |  |  |  |  |
| (Intercept) | 5.964 | 2.277 | 2.620 | **0.009** |
| Log(Strongyles EPG + 1) | -1.439 | 0.565 | -2.548 | **0.011** |
| Log(*E. Cervi* LPG + 1) | 0.247 | 0.276 | 0.896 | 0.370 |
| Log(*F. Hepatica* EPG + 1) | -0.839 | 0.365 | -2.297 | **0.022** |
| Sex [male] | 0.746 | 0.817 | 0.914 | 0.361 |
| Deer year [2017] | 0.319 | 0.845 | 0.377 | 0.706 |
| Deer year [2018] | 0.576 | 1.231 | 0.468 | 0.640 |
| **Model E (Autumn, yearling survival, n=65)** |  |  |  |  |
| (Intercept) | 0.801 | 1.425 | 0.562 | 0.574 |
| Log(Strongyles EPG + 1) | -1.884 | 0.698 | -2.697 | **0.007** |
| Log(*E. Cervi* LPG + 1) | -0.046 | 0.258 | -0.176 | 0.860 |
| Log(*F. Hepatica* EPG + 1) | 0.451 | 0.333 | 1.353 | 0.176 |
| Sex [male] | -0.457 | 0.756 | -0.605 | 0.545 |
| Deer year [2017] | 3.564 | 1.662 | 2.145 | **0.032** |
| Deer year [2018] | 3.394 | 1.999 | 1.698 | 0.090 |

**Figure 1:**
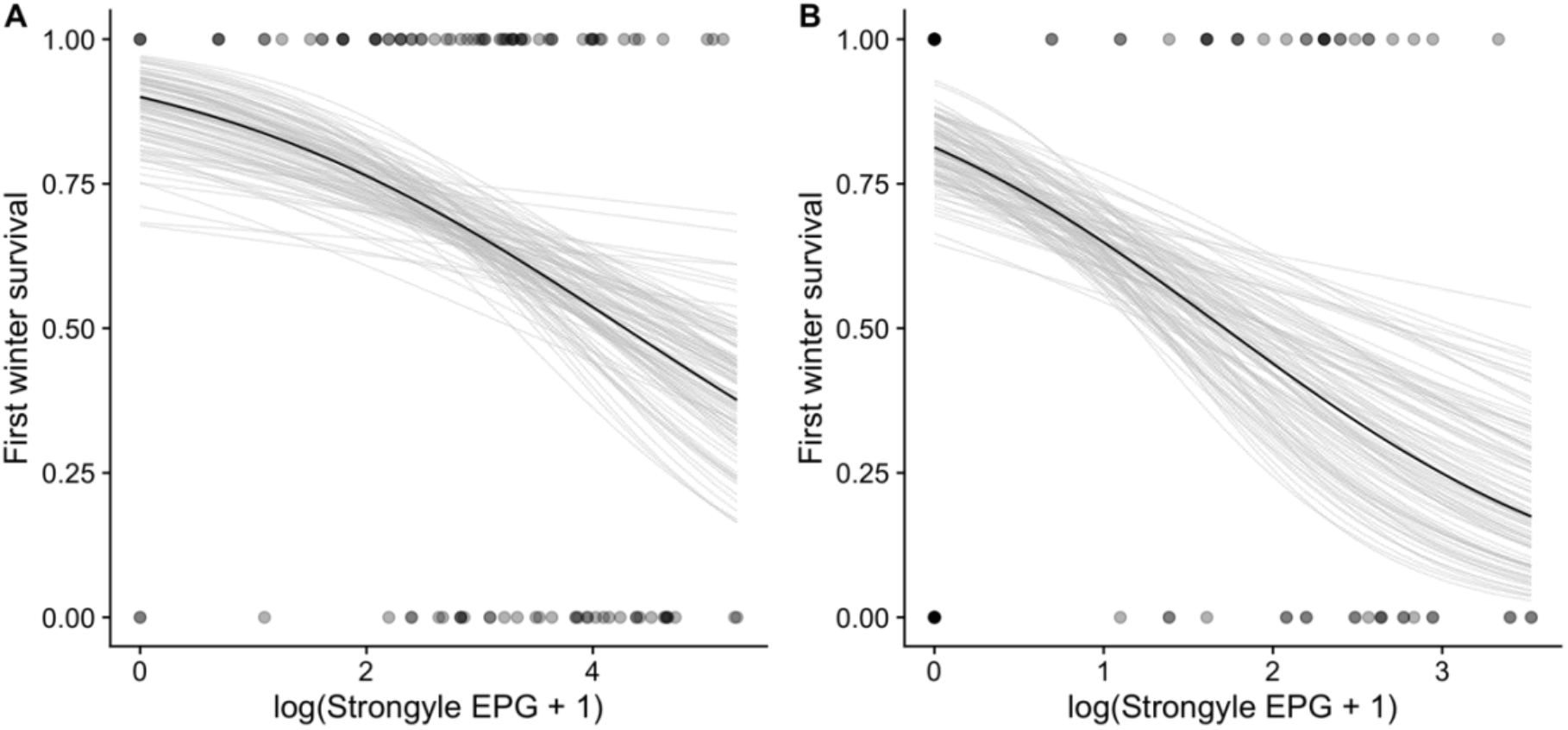
Probability of calf survival over their first winter (1=survived, 0=died) as predicted by their strongyle FEC (log(EPG+1)) from samples taken in the (A) summer (model A) and (B) autumn (model B). Solid black line = fitted logistic regression slope. Transparent grey lines = 100 random draws from model estimates to display variation in the estimated slope. Transparent grey dots = individual sample.

An individual’s spring strongyle FEC was significantly negatively associated with survival over their second winter as yearlings (model C; -0.869 ±0. 372, p=0.019). Individuals with the lowest spring strongyle FECs (<10 EPG, 13.4% of samples) had a >95.4% probability of survival over their second winter as a yearling, and those with the highest spring strongyle FECs (>90 EPG, 26.9% of samples) had a <76.8% probability of survival (Figure 2A). A yearling’s summer strongyle FEC was also significantly negatively associated with their overwinter survival (model D; -1.44 ±0.565, p=0.011). Yearlings with the lowest summer strongyle FECs (<5 EPG, 10.9% of samples) had a >96.2% probability of survival over their second winter, whilst those with the highest summer strongyle FECs (>30 EPG, 18.75% of samples) had a <70.6% probability of survival (Figure 2B). A yearling’s autumn strongyle FEC was significantly negatively associated with survival over their second winter (model E; -1.88 ±0.698, p=0.007). Yearlings with the lowest autumn strongyle FECs (0 EPG, 53.8% of samples) had a 97.1% probability of overwinter survival, whilst those with the highest autumn strongyle FECs (10 EPG, 12.3% of samples) had a <26.7% probability of survival (Figure 2C). A deer’s *F. hepatica* FEC was negatively associated with subsequent overwinter survival only in yearling summer samples (model D; -0.839 ±0.365, p=0.022). Yearlings with the lowest summer *F. hepatica* FECs (0 EPG, 18.8% of samples) had a 97.4% probability of survival over their second winter, whilst those with the highest summer *F. hepatica* FECs (>30 EPG, 18.8% of samples) had a <67.5% probability of survival (Figure 2D). A deer’s *E. cervi* FEC was not significantly associated with subsequent overwinter survival in any sampled season (Table 2). In this sample of deer there was also no significant difference in calf or yearling overwinter survival between males and females (Table 2). Calf and yearling overwinter survival varied between years in models using data sampled from autumn, with lower survival probabilities in 2016 compared to 2017 and 2018 (model B and model E, Table 2).

**Figure 2:**
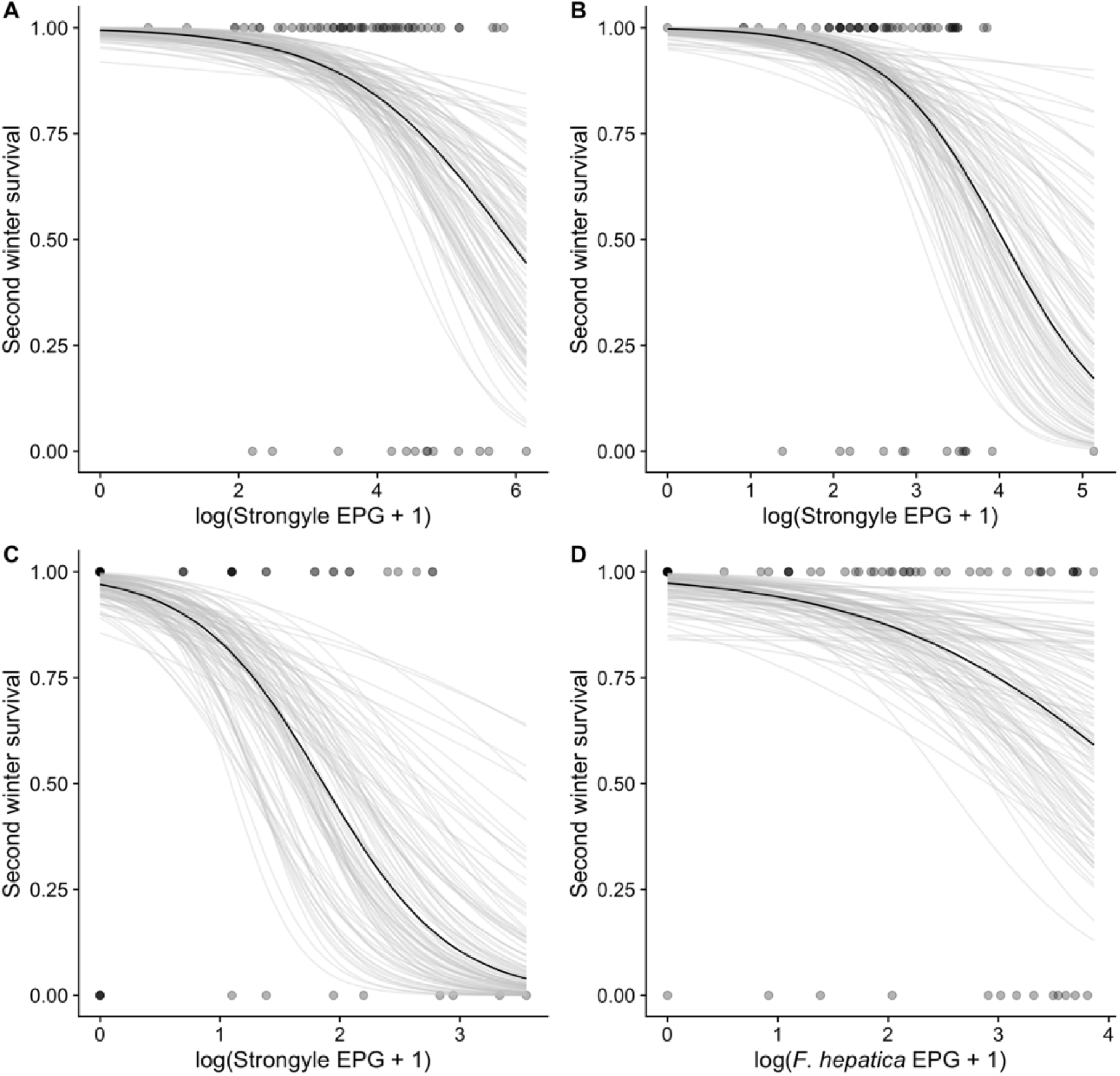
Probability of yearling survival over their second winter (1=survived, 0=died) as predicted by their strongyle FEC (log(EPG+1)) from samples taken in the (A) spring as calves (model C), (B) summer as yearlings (model D) and (C) autumn as yearlings (model E). And as predicted by (D) *F. hepatica* FEC (log(EPG+1)) from samples taken in the summer as yearlings (model D). Solid black line = fitted logistic regression slope. Transparent grey lines = 100 random draws from model estimates to display variation in the slope. Transparent grey dots = individual samples.

## 4. Discussion

We provide observational evidence that parasite infection is associated with substantially reduced survival probability in young red deer. Individuals with higher strongyle nematode intensities showed a reduced overwinter survival probability, consistent with the observation that strongyle infection is negatively correlated with fitness in adult females (Albery *et al*., 2021). While our analysis cannot infer causality, adult strongyle nematodes are known to cause damage to their hosts’ abomasal mucosa, and consequently cause disruption to nutrient absorption in ungulates (Hoberg *et al*., 2001). Indeed, studies experimentally removing helminths by administration of anthelminthic treatment have shown strongyle nematodes to cause mortality in other wild mammals e.g. Soay sheep (*Ovis aries*) (Coltman *et al*., 1999; Gulland, 1992), reindeer (*Rangifer tarandus*) (Albon *et al*, 2002), and snowshoe hares (*Lepus americanus*) (Murray *et al*., 1997). Taking this evidence together, it is therefore reasonable to consider that strongyle nematodes are having negative impacts on the health of juvenile red deer and are contributing towards overwinter mortality.

Studies of juvenile Soay sheep have uncovered a negative effect of strongyle nematodes on survival, in addition to the effects of body weight, a correlate of body size (Sparks *et al*., 2020). A similar effect may be occurring in red deer, but development of a non-invasive measure of body size for the Rum study system would be necessary to disentangle size- and parasite-dependent effects on survival. Nonetheless, our analysis shows a survival cost associated with strongyle infection in juvenile red deer which may exert positive selection on resistance to infection, as has been observed in other ungulate study systems (Hayward *et al*., 2011). Furthermore, this negative association is observed despite low mean strongyle egg counts in both calves and yearlings compared to the mean strongyle counts that are observed in lamb and yearling Soay sheep (Craig *et al*., 2008). Strongyle egg counts peaked in calves sampled in spring (April), which may reflect a transmission strategy of coinciding maximum propagule output with the influx of immunologically naïve calves in May. The low intensities and prevalence of strongyle eggs in the autumn is likely due to a reduction in propagule output, as colder temperatures decrease transmission, rather than reductions in actual burden (Albery *et al*., 2018). In general, calves had higher strongyle intensities than yearlings, agreeing with previous findings, which may result from the negative effects of strongyle infection on juvenile overwinter survival and/or the maturation of the naïve immune system (Albery *et al*., 2018).

In the case of *F. hepatica*, yearling overwinter survival was predicted by the individual’s count in summer, but not by its count in spring or autumn. This may be true seasonal variation, or a result of the selection of samples used in the yearling analysis (models C, D, and E). Only ∼16% of yearlings died in their second winter (in contrast to ∼37% of calves in their first year), which may have reduced the models’ ability to reliably detect an association between *F. hepatica* FECs and survival. Ultimately, the analysis presented is restricted in estimating the association between *F. hepatica* and juvenile survival; collection of further *F. hepatica* FECs and fitness data from yearlings will be necessary to better understand their survival effects. *F. hepatica* is known to have a negative effect on weight gain in domestic cattle and sheep (Hayward *et al*., 2021). Similar effects of *F. hepatica* infection in wild red deer may explain their association with a reduced survival probability, as lighter individuals are less able to survive over winter periods of poor nutrition (Loison *et al*., 1999).

In contrast to strongyles and *F. hepatica, E. cervi* did not have any apparent survival effects. *E. cervi* nematodes infect the central nervous system and skeletal muscles of their hosts and propagated larvae migrate through the bloodstream to the lungs prior to being swallowed and excreted (Mason, 1989). Descriptions of the clinical symptoms of disease from *E. cervi* infection have included paresis of hind limbs and pneumonia; however, pathogenicity is relatively low in red deer in Scotland (Mason, 1989). Minimal pathology of *E. cervi* infection in juvenile red deer may explain its lack of association with subsequent overwinter survival probability. Furthermore, this result may reflect a host response of tolerance to *E. cervi* infection, where minimising the damage caused by infection is prioritised over eradicating the worms (McSorley and Maizels, 2012). This strategy could explain the high intensities and prevalence, and lack of age-bias of this parasite in the population (Albery *et al*., 2018). There was also no sex disparity in survival probability, contrary to expected male-biased mortality (Moore & Wilson, 2002). However, this observation is not likely due to sex differences in parasite FECs, which were small in calves and yearlings for strongyles, and not observed for *F. hepatica* (Albery *et al*., 2018).

Host population density is predicted to positively affect helminth transmission (Tompkins *et al*., 2001); at higher population densities juveniles may show higher intensities of helminth infection, as has been observed in another wild ungulate population, Soay sheep (Hayward *et al*., 2014). Considering the survival costs of strongyle infection in juveniles demonstrated here, density-dependent parasitism could be involved in the density-dependent juvenile survival that occurs in the deer (Coulson *et al*., 1997). There is limited understanding of how parasites may regulate ungulate hosts populations; however, experimental studies of wild reindeer suggest helminths may be capable of regulating the population via density-dependent effects on host reproduction (Albon *et al*., 2002). Strongyle nematodes are likely to have mediating effects on population dynamics in red deer, by reducing juvenile survival and by reducing survival and future reproduction in adult females (Albery *et al*., 2021). Whilst the observational nature of the Rum red deer study system precludes the manipulation of helminth infection necessary to determine a regulatory role, collection of further years of parasite and fitness data paired with population density data would be valuable in developing a more nuanced understanding of how helminths impact wild populations. Furthermore, additional years of longitudinal parasite and fitness data collection will inform the long-term effects of juvenile parasitism on future fitness as deer are studied through to maturity and senescence.

## Acknowledgements

Field data collection was supported by the UK Natural Environment Research Council (core grant and PhD studentship to GFA). GFA is currently supported by a College for Life Sciences Fellowship from the Wissenschaftskolleg zu Berlin. We are grateful to NatureScot for permission to work on Rum.

